# Species hybridisation and clonal expansion as a new fungicide resistance evolutionary mechanism in *Pyrenophora teres* spp

**DOI:** 10.1101/2021.07.30.454422

**Authors:** Chala Turo, Wesley Mair, Anke Martin, Simon Ellwood, Richard Oliver, Francisco Lopez-Ruiz

## Abstract

The barley net blotch diseases are caused by two fungal species of the *Pyrenophora* genus. Specifically, spot form net blotch is caused by *P. teres* f. sp. *maculata* (*Ptm*) whereas net form net blotch is caused by *P. teres* f. sp. *teres* (*Ptt)*. *Ptt* and *Ptm* show high genetic diversity in the field due to intraspecific sexual recombination and hybridisation of the two species although the latter is considered rare. Here we present occurrence of a natural *Ptt/Ptm* hybrid with azole fungicides resistance and its implication to barley disease management in Australia. We collected and sequenced a hybrid, 3 *Ptm* and 10 *Ptt* isolates and performed recombination analyses in the intergenic and whole genome level. Eleven out of 12 chromosomes showed significant (*P <* 0.05) recombination events in the intergenic regions while variable recombination rate showed significant recombination across all the chromosomes. Locus specific analyses of *Cyp51A1* gene showed at least four recombination breakpoints including a point mutation that alter target protein function. This point mutation did not found in *Ptt* and *Ptm* collected prior to 2013 and 2017, respectively. Further genotyping of fourteen *Ptt,* 48 HR *Ptm*, fifteen *Ptm* and two *P. teres* isolates from barley grass using Diversity Arrays Technology markers showed that all HR *Ptm* isolates were clonal and not clustered with *Ptt* or *Ptm*. The result confirms occurrence of natural recombination between *Ptt* and *Ptm* in Western Australia and the HR *Ptm* is likely acquired azole fungicide resistance through recombination and underwent recent rapid selective sweep likely within the last decade. The use of available fungicide resistance management tactics are essential to minimise and restrict further dissemination of these adaptive HR *Ptm* isolates.

## INTRODUCTION

Spot form net blotch (SFNB) caused by *Pyrenophora teres* f. sp. *maculata* (Smedegard-Petersen, 1971) (*Ptm*) and net form net blotch (NFNB) caused by *Pyrenophora teres* f. sp. *teres* (anamorph *Drechslera teres* [Sacc.] Shoem.) (*Ptt*) are two closely related major fungal pathogens of barley (*Hordeum vulgare* L.). Worldwide importance of the two forms varies, with one or the other predominant in a particular region based on environmental, agronomic, and varietal factors (McLean *et al.*, 2009). The pathogens can cause yield losses of 10 - 44% (Steffenson *et al.*, 1991; Jebbouj & El Yousfi, 2009; McLean *et al.*, 2009). In Australia, the damage inflicted by both pathogens has been estimated to be on average AUD$62 million with potential losses estimated at AUD$309 million annually in the absence of control measures (Murray & Brennan, 2010). Symptoms induced by both pathogens can be easily distinguished, with *Ptm* producing dark brown spots and *Ptt* dark net-like lesions along the leaves (Smedegard-Petersen, 1971).

Demethylase-inhibitor (DMI) fungicides are one of the primary modes of action (MOA) employed as part of the chemical management of SFNB and NFNB (Akhavan *et al.*, 2017). However, the widespread use of DMI chemistry has led to the emergence of fungicide resistance in recent years. Mutation F489L in the DMI fungicide target gene, *Cyp51A*, was found to be associated with DMI resistance in *Ptt* isolates since 2013 in Western Australia (WA)(Mair *et al.*, 2016). From 2016 onwards, *Ptm* isolates resistant to DMI fungicides were found to carry mutation F489L and/or an insertion of a 134 bp fragment of a transposable element in the *Cyp51A* promoter region (Mair *et al.*, 2020). *Cyp51A* exists as a single copy in *Ptm* while two copies have been identified in *Ptt* (Mair *et al.*, 2016).

Worldwide studies on population genetics of *Ptt* and *Ptm* have shown extremely high levels of genetic diversity in field populations (Rau *et al.*, 2003; Serenius *et al.*, 2007; Bogacki *et al.*, 2010; McLean *et al.*, 2010; Gupta *et al.*, 2012; McLean *et al.*, 2014; Akhavan *et al.*, 2016). These genetic variations were driven by sexual reproduction, gene flow and transposable element invasion. More recently, *Ptt* population genetic studies using Diversity Arrays Technology (DArT) found rapid changes in population genetic structure under field conditions (Poudel *et al.*, 2019b).

Continuous growth of barley can lead to build up of both *Ptt* and *Ptm* populations, which favours their genetic diversification and ultimately the emergence of new pathotypes through sexual recombination. Recombination can rapidly assemble new combinations of alleles to overcome selection pressure, invade novel niches or lead to emergence of new pathogenic species (McDonald & Linde, 2002; Depotter *et al.*, 2016).

In addition to intraspecific sexual recombination, occurrence of interspecific hybridization between *Ptt* and *Ptm* has been reported by several authors. The first putative hybrid was reported from South Africa in 2002 (Campbell *et al.*, 2002), followed by the Czech Republic (Leisova *et al.*, 2005) and Australia (Lehmensiek *et al.*, 2010; McLean *et al.*, 2014). These recombinant hybrids are able to retain their genetic stability over time (Campbell & Crous, 2003; Jalli, 2011)). The symptoms induced by hybrids are indistinguishable from those produced by *Ptt* or *Ptm*, and some hybrids show more virulence than either parents (Campbell *et al.*, 1999; Campbell *et al.*, 2002; Campbell & Crous, 2003).

Interspecific hybridization in *P. teres* have previously been studied using amplified fragment length polymorphisms (AFLP), random amplified polymorphic DNA (RAPD) or form-specific PCR markers, which depend on comparatively small number of molecular markers (Campbell *et al.*, 2002; Rau *et al.*, 2003; Lehmensiek *et al.*, 2010; McLean *et al.*, 2014; Poudel *et al.*, 2017). Population genetic analyses can now be performed using higher throughput technologies such as DArT (Ndjiondjop *et al.*, 2017). Furthermore, the ever decreasing costs of whole genome sequencing allowed population genomic studies to uncover the level of introgression occurring across diverse microbes. Such studies were applied to detect recombination among strains or closely related species of *Z*y*moseptoria* species (Stukenbrock & Dutheil, 2017), *Aspergillus fumigatus* (Abdolrasouli *et al.*, 2015), *Blumeria* species (Feurtey *et al.*, 2019) and *Histoplasma* species (Maxwell *et al.*, 2018). In some instances, introgressed regions have been associated with responses to stresses (Zhang *et al.*, 2018) or adaptation to new hosts (Feurtey *et al.*, 2019).

The almost simultaneous emergence of the same DMI resistance mutation in both *Ptt* and in highly resistant *P. teres* isolates that induced typical spot form symptoms (termed here “HR *Ptm*”), raises the question as to whether fungicide resistance emerged as a result of hybridization between these two pathogens. Here we present a comprehensive genetics and genomic studies that dissects the occurrence of natural hybridization between *Ptt* and *Ptm*, and determines the origin of fungicide resistant hybrids.

## MATERIALS AND METHODS

### Isolate collection and species identification using form-specific primers

A total of 48 HR *Ptm* isolates were obtained from diseased leaf and stubble samples collected in the 2017 and 2018 cropping seasons from nine locations across the barley growing regions of WA (Table 1). Fungal isolations from infected leaves were conducted as previously described (Mair *et al.*, 2020). For isolation from infected stubble, tissue was washed with sterile water, dried on paper towel, placed in petri dishes on moist paper, and incubated at 15 °C for 5 days. A single conidium was transferred to V8PDA (10 g potato-dextrose agar, 3 g CaCO_3_, 15 g agar, 150 mL V8 juice in 850 mL deionized H_2_O) plates amended with ampicillin (100 μg mL^−1^), streptomycin (100 μg mL^−1^), and neomycin (50 μg mL^−1^). The isolates were grown on V8PDA for 5 days at room temperature. Genomic DNA was isolated from mycelia on a Biosprint 15 instrument using Biosprint DNA Plant Kit (QIAGEN, Hilden, Germany) as per manufacturer’s instructions. Genomic DNA was visualized by electrophoresis on 1% agarose gel, assessed for purity on a Nanodrop 2000 (Thermo Fisher Scientific, Waltham, MA USA), and quantitated using a Quantus fluoromoter with the QuantiFluor dsDNA kit (Promega, Madison, WI, USA). The isolates were identified using form-specific molecular markers and PCR conditions described in Poudel *et al.* (2017).

**Table 1.**
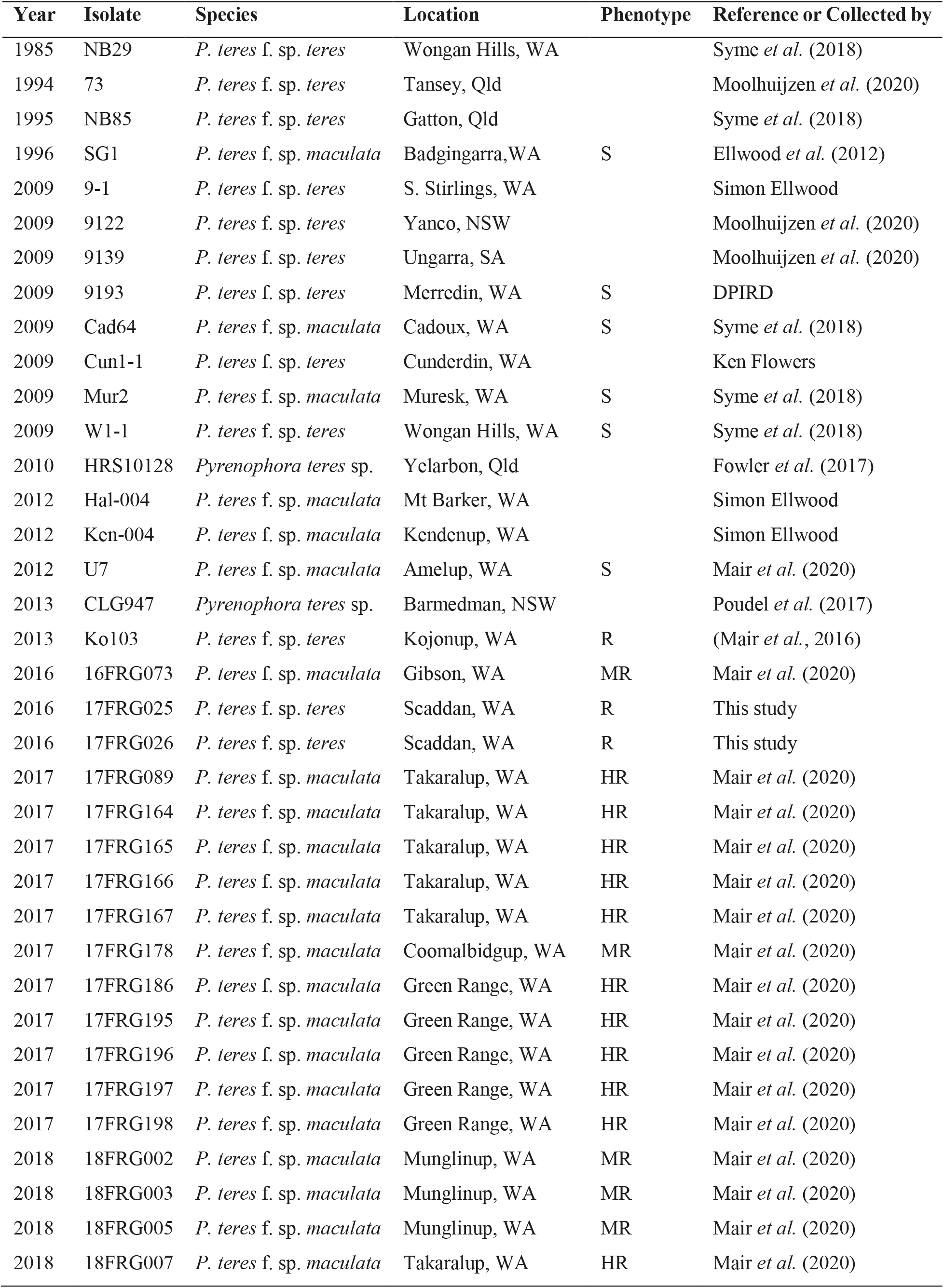
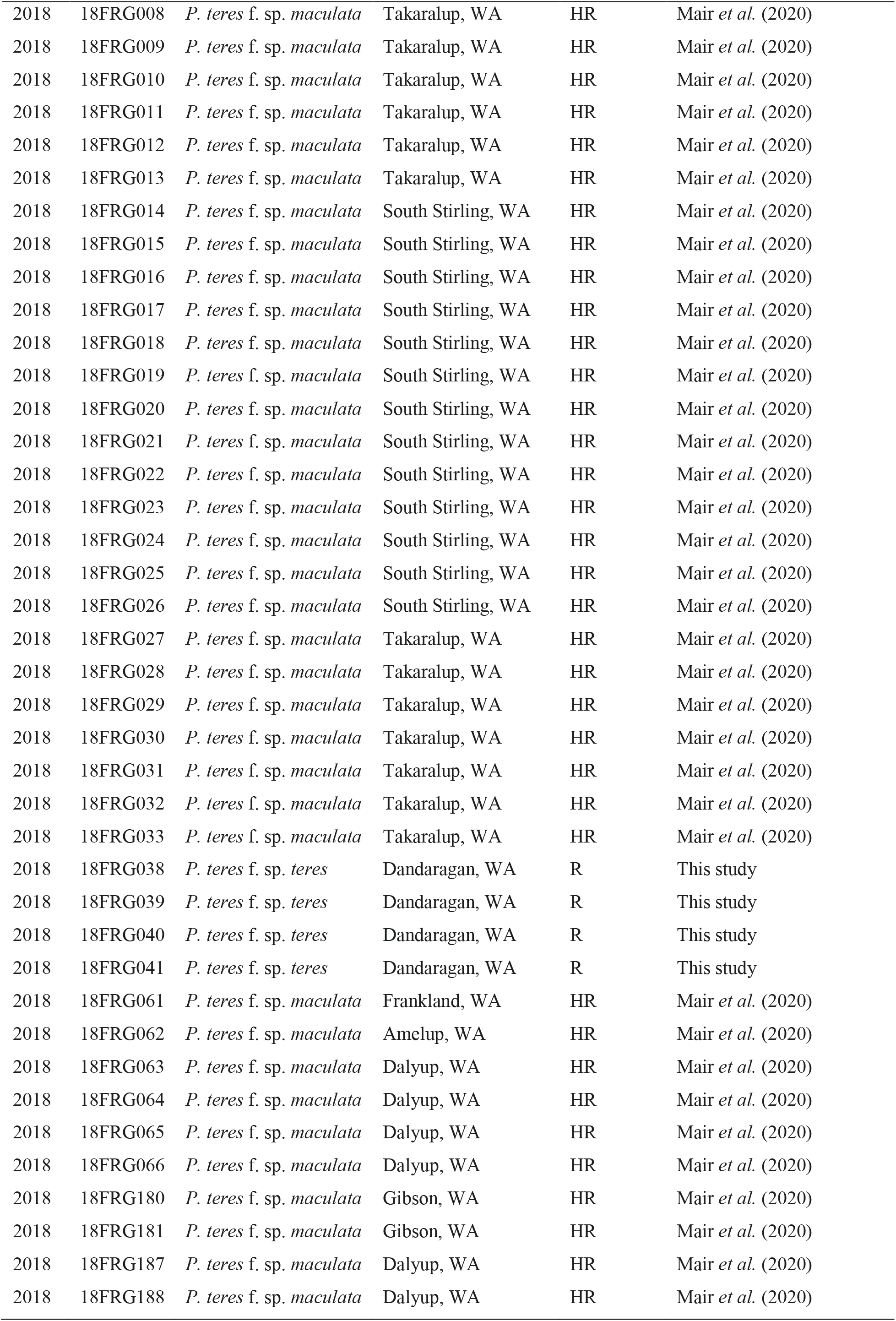
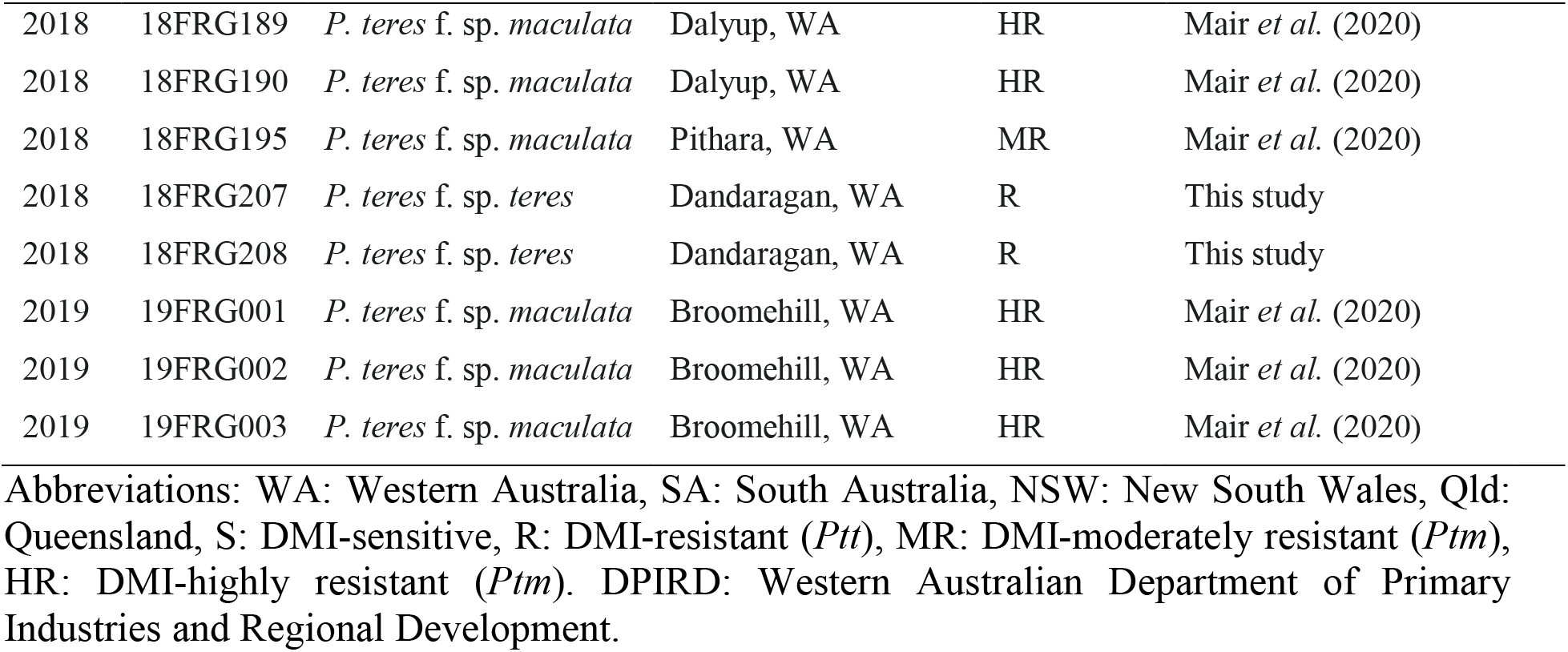
Description of isolates used in the current study

### Whole genome sequencing and assembling

For whole genome sequencing, isolates 9193, Ko103, 16FRG073, 17FRG026, and 17FRG089 were cultured and genomic DNA extracted as described by Syme *et al.* (2018). DNA was assessed for purity and quantitated as described above. Sequencing was performed by Macrogen (Seoul, South Korea) using 20 kbp SMRTbell libraries on the PacBio RSII platform (Pacific BioSciences, Menlo Park, CA, USA) as per manufacturer’s protocols. The reads were assembled into contigs using canu version 1.6 (Koren *et al.*, 2017).

### Genetic diversity analysis

SilicoDArT and SNP DArT markers were used to analyse the genetic diversity of 48 DMI resistant isolates collected from WA. 15 *Ptm*, 14 *Ptt* isolates collected between 1996 and 2019 together with two *P. teres* isolates obtained from barley grass grass (*Hordeum leporinum*) were also included in the study. DArTseq was performed by Diversity Arrays Technology Pty Ltd (Canberra, ACT, Australia) with the *P. teres* array using the HiSeq 2000 platform (Illumina, San Diego, CA, USA) as per manufacturer’s protocols. Only markers with sequences present in the W1-1 (GenBank BioProject PRJEB18107, assembly GCA_900232045.2) or the 0-1 (GenBank BioProject PRJNA392275, assembly GCA_006112615.1) reference genomes were used, and markers with more than 5% missing data were removed; the remaining 9,656 silicoDArT and SNP markers were used in the analysis. The similarity matrix was constructed using the DICE coefficient (Dice, 1945) in the Dissimilarity program of the Dissimilarity Analyses and Representation for Windows (DARwin) software package, version 6.0.21 (http://darwin.cirad.fr/) (Perrier & Jacquemoud-Collet, 2006). Cluster analyses of the matrix values was performed using the Unweighted Pair-Group Method with Arithmetic Mean (UPGMA) (Sneath & Sokal, 1973) in the Hierarchical Clustering program, and the dendrogram was produced using Trees Draw program of DARwin.

DArT SNP markers generated for each fungal species were used to compute genetic diversity using R statistical software (https://www.r-project.org/)(R Core Team, 2018). The raw DArT data were imported and converted into genlight object using gl.read.dart() function implemented in dartR package (Gruber *et al.*, 2018). Fixation indexes (*F_ST_*) were computed using ‘stamppFst’ to explore genetic differentiation, as implemented in R statistical package ‘StAMPP’ (Pembleton *et al.*, 2013). The genetic similarity of individuals was analysed and visualized using Principal Coordinates Analyses (PCoA) using dartR statistical package (Gruber *et al.*, 2018) developed for analyses of DArT markers. The PCoA was based on standardized covariance of genetic distances calculated from DArT markers and the analyses was performed using default parameters. The Nei’s genetic distances between pairs were computed using the StAMPP package (Pembleton *et al.*, 2013).

### Analyses of recombination in the intergenic regions

The reference genome sequence of *Ptt* isolate W1-1 (Syme *et al.* (2018) was used in this analysis. Coordinates of protein coding genes were taken from the General Feature Format (GFF) file from W1-1, after adding 500 bp upstream and downstream of each gene using the slope command available in the Bedtools suite (Quinlan & Hall, 2010). Any regions that overlapped with repetitive elements were discarded and the remaining coordinates were used to extract 1 kb intergenic regions. Homologous regions were searched from other *Ptt* and *Ptm* genomes sequenced in this study (9193, Ko103, 16FRG073, 17FRG026, and 17FRG089), as well as the previously published genomes of isolates SG1 (Ellwood *et al.*, 2012), NB29, NB85, Cad64 and Mur2 (Syme *et al.*, 2018), and 73, 9122 and 9139 (Moolhuijzen *et al.*, 2020) using Basic Local Alignment Search Tool (BLAST) (Altschul *et al.*, 1990), and the best matching regions were extracted to generate multiple sequence alignments for each intergenic regions using MAFFT (Katoh, 2002). The aligned regions were subjected to recombination test using PHIPack (Bruen, 2005).

### Whole genome sequence based recombination analysis

The genome sequences of nine *Ptt* isolates (two DMI-resistant: Ko103 and 17FRG026, and seven DMI-sensitive: 73, 9122, 9139, 9193, NB29, NB85 and W1-1), four *Ptm* isolates (one DMI-resistant: 16FRG073, and three DMI-sensitive: Cad64, Mur2 and SG1), and one DMI-resistant putative hybrid (“HR *Ptm*”, 17FRG089), were investigated to determine whether recombination events between *Ptt* and *Ptm* isolates had occurred. Genome sequences were aligned using Sibeliaz (Minkin & Medvedev, 2019) using sequence of eleven k-mers. The multiple sequence alignments were filtered (MinBlockLength of 100 bp), duplicates removed, concatenated and ordered relative to W1-1 after masking any repetitive regions. The ordered alignments were filtered using MafFilter (Dutheil *et al.*, 2014). Only blocks where all isolates were present were retained; a window of 10 bp was slided by 1 bp, and windows containing at least two insertion and or deletion events were discarded and the containing blocks split; in addition, windows with a total of > 100 gap characters were discarded and the containing blocks were split; and all blocks were merged according to reference genome with empty positions filled by “N”. Refined multiple sequence alignments were subjected to variable recombination rate analyses using LDJump (Hermann *et al.*, 2019) with segment length of 5 kb interval, *P* value of 0.05. LDJump requires other recombination analyses tools including LDhat (Auton & McVean, 2007) and PhiPack to estimate variable recombination rate and generate a recombination map in a specific DNA segment. Final recombination maps were generated for each chromosome using LDJump. The segment length was set above recommended 1 kb to avoid imputation due to low SNP density.

### Phylogenetic analysis

Phylogenetic analyses was performed on selected intergenic regions and all the twelve chromosomes. In intergenic regions, multiple sequence alignment was performed using MAFFT version 5 (Katoh, 2002) with default parameters. Phylogenetic trees were generated for selected genomic regions following the neighbour-joining method with Juke-Cantor genetic distance with bootstrap sampling of 1,000 random replicate sampling and 1,000,000 Seeding in Genious version 8.0.3 (Kearse *et al.*, 2012). Similarly, in chromosome level sequence alignments, the phylogenetic trees were constructed using modified neighbour-joining method (Gascuel, 1997) using maximum likelihood distance estimated with pairwise substitution model K80 (kappa=2) as implemented in MafFilter program and trees were visualized using FigTree V1.4.4 (http://tree.bio.ed.ac.uk/software/figtree/).

## RESULTS

### Fungicide resistant isolates show evidence of hybridisation between *Ptt* and *Ptm* and recombination of the mutant *Cyp51A* allele

Independent repeated PCR analyses using *Ptm*- and *Ptt*- specific primers showed all HR *Ptm* isolates were positive for all 6 *Ptm* specific markers (Figure 1). Five of the *Ptt*-specific markers amplified only all of the *Ptt* isolates, while one *Ptt*-specific marker (PttQ4) was able to amplify both all *Ptt* and all HR *Ptm* isolates. On the other hand, all six of the *Ptm*-specific markers amplified only *Ptm*. An identical pattern was observed across all 48 HR *Ptm* isolates screened using the form-specific markers.

**Figure 1.**
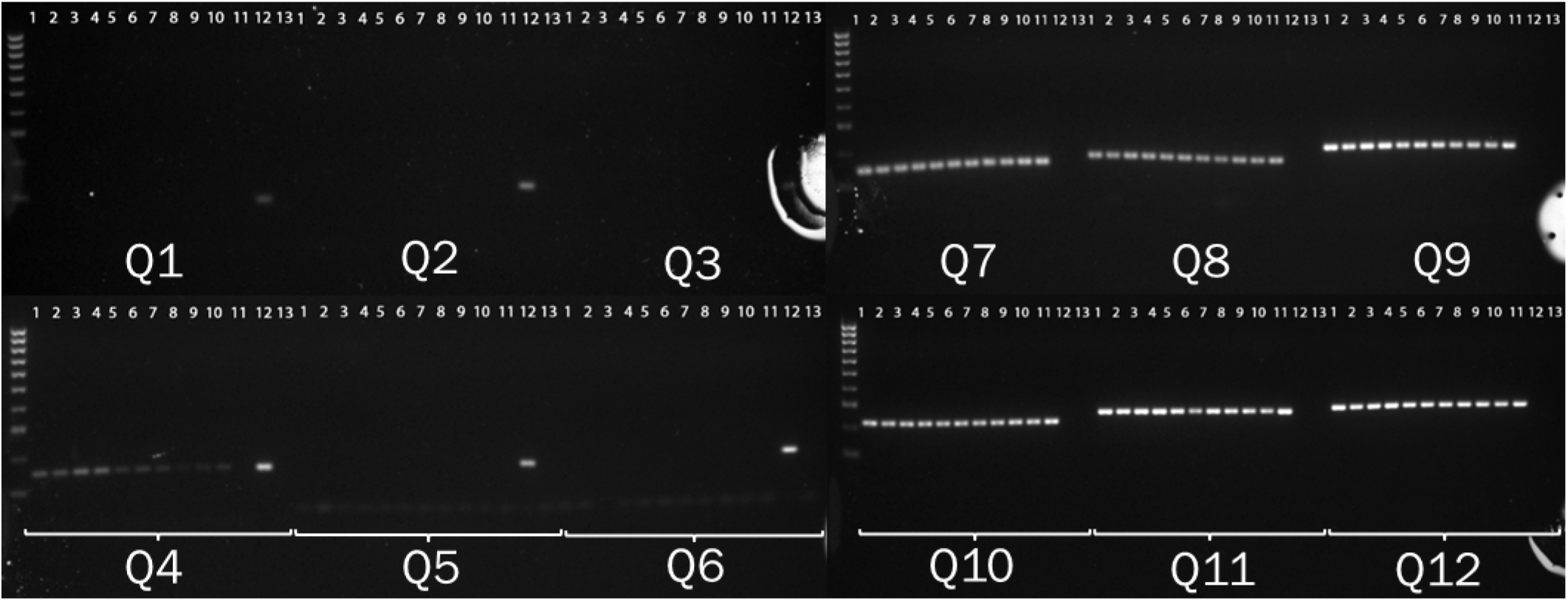
Amplification of using form specific markers. The lanes 1. 17FRG164 2. 17FRG165 3. 17FRG166 4. 17FRG167 5. 17FRG186 6. 17FRG187 7. 17FRG195 8. 17FRG196 9. 17FRG197 10. 17FRG198 11. SG1 (*Ptm* Control) 12. Ko103 (*Ptt* Control) 13. No Template Control. Lane 1-10 isolates carry F489L (c1467a) mutation in highly resistant *Ptm* isolates.

The multiple sequence alignments of 64 *Cyp51A* coding sequences from *Ptt* and *Ptm* indicated a preferential mutation pattern in all the HR *Ptm* isolates. All HR *Ptm* isolates carry the SNP c1467a for the amino acid substitution F489L, identical to the SNP found in all DMI resistant *Ptt* (Figure 2). At the proximal polymorphic site in the *Cyp51A* sequence (position 1488), all HR *Ptm*, as well as all *Ptt* (both DMI-sensitive and resistant) carry variant g1488; all other *Ptm* (both DMI-sensitive and resistant) carry c1488. Genetic recombination rate analyses using DnaSP identified at least four break within *Cyp51A* (Figure 2).

**Figure 2.**
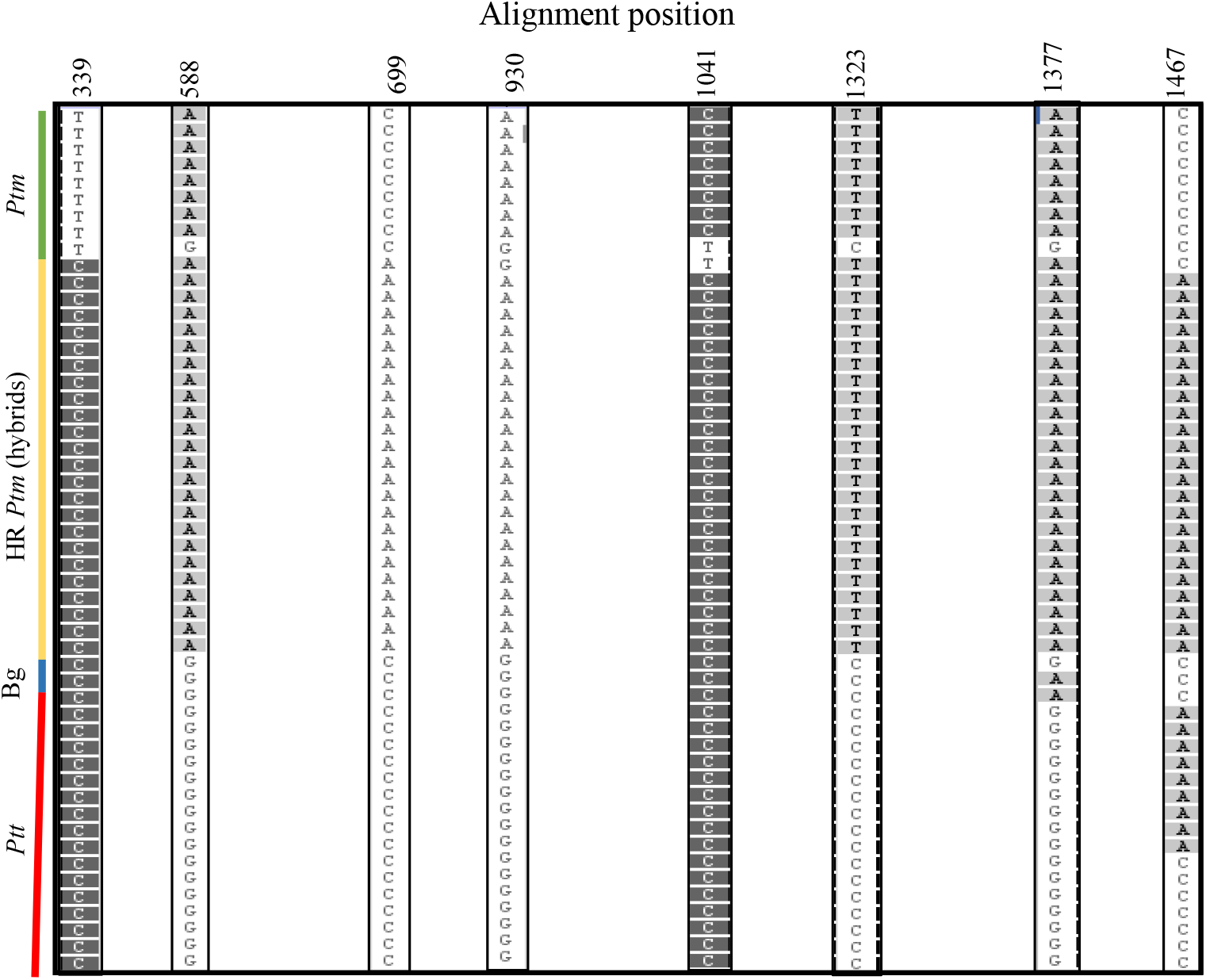
The four recombination break points predicted in CYP51A protein coding region identified using DnaSP. Position 1467 is a mutation (c1467a) that alter the protein function and observed only in HR *Ptm* and *Ptt.*

### Hybrid genome shows evidence of hybridization between Ptt and Ptm

We further tested signatures of hybridization among isolates of *Ptt, Ptm* and HR *Ptm* first in the conserved intergenic regions and then extended to chromosome level. A total of 1,393 intergenic loci with 1 kb length were identified and tested for presence of recombination on the twelve *Ptt* chromosomes and their respective homologs in *Ptm* and HR *Ptm.* The recombination analyses using PhiPack revealed the existence of significant (*P < 0.05*) recombination events on 75 of the loci (Table 2). The number of significant recombination events across intergenic regions ranged from twelve on Chr02 to two on Chr04, while no significant intergenic recombination was detected on Chr12.

**Table 2.**
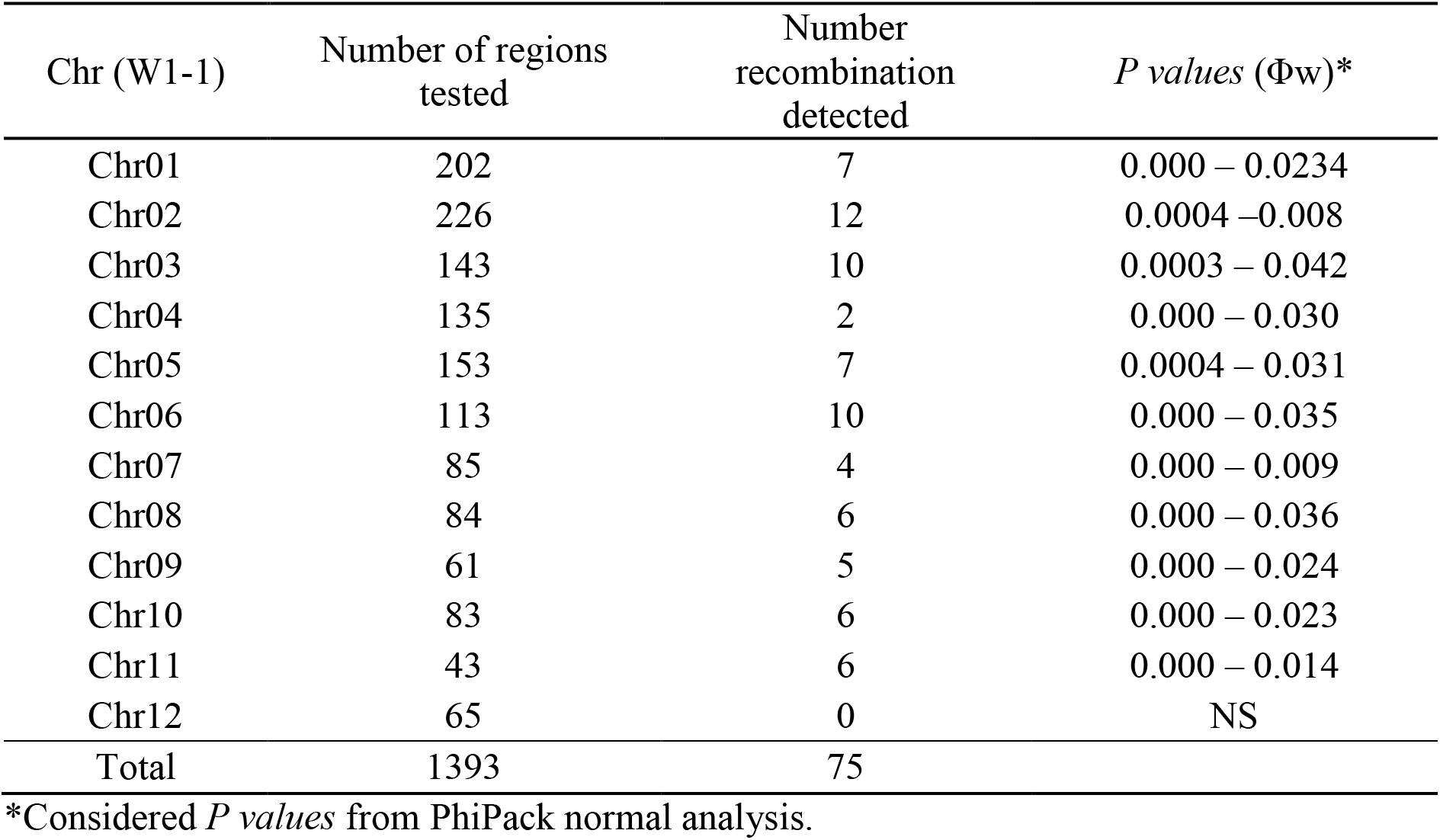
Recombination frequency detected in the intergenic 1 kb regions across 12 chromosomes

| Chr (W1-1) | Number of regions tested | Number recombination detected | *P* values ($\Phi_w$)* |
| --- | --- | --- | --- |
| Chr01 | 202 | 7 | 0.000 – 0.0234 |
| Chr02 | 226 | 12 | 0.0004 – 0.008 |
| Chr03 | 143 | 10 | 0.0003 – 0.042 |
| Chr04 | 135 | 2 | 0.000 – 0.030 |
| Chr05 | 153 | 7 | 0.0004 – 0.031 |
| Chr06 | 113 | 10 | 0.000 – 0.035 |
| Chr07 | 85 | 4 | 0.000 – 0.009 |
| Chr08 | 84 | 6 | 0.000 – 0.036 |
| Chr09 | 61 | 5 | 0.000 – 0.024 |
| Chr10 | 83 | 6 | 0.000 – 0.023 |
| Chr11 | 43 | 6 | 0.000 – 0.014 |
| Chr12 | 65 | 0 | NS |
| Total | 1393 | 75 |  |

\*Considered *P* values from PhiPack normal analysis.

For further large-scale recombination analysis, we generated multiple sequence alignments varying between 3.84 Mb on Chr01 and 0.94 Mb on Chr12. Occurrence of recombination was observed on all the twelve chromosomes. Chr11 had the highest recombination rate (0.0437) followed by Chr06 (0.0394) (Figure 4). As opposed to the intergenic regions, all twelve of the chromosomes showed significant evidence of recombination, which appeared to occur in clusters across certain chromosomal regions as indicated on the recombination map shown in figure 4.

**Figure 3.**
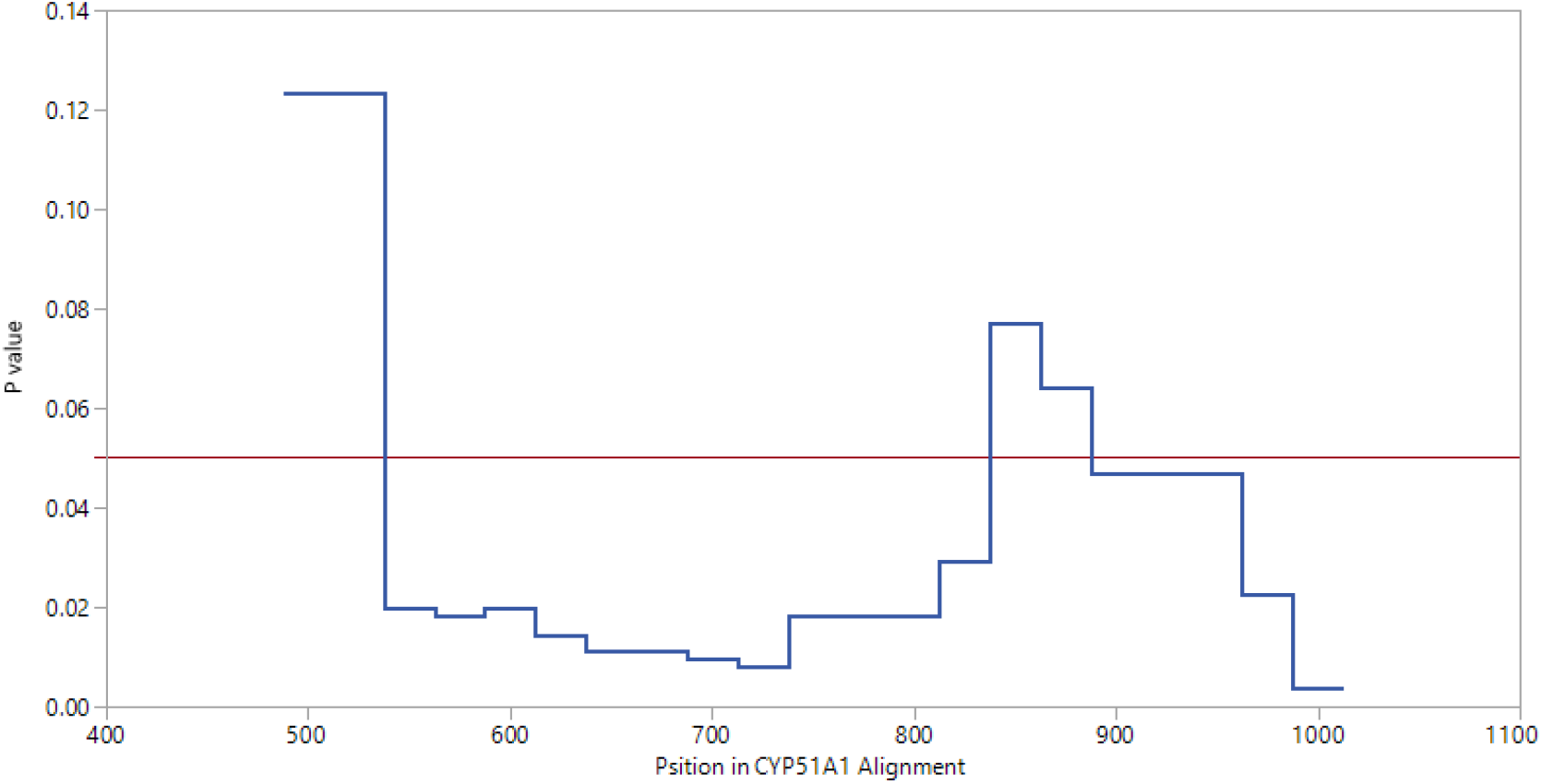
Potential recombination regions predicted in CYP51A protein coding sequences. The red horizontal line is drawn at *P value* of 0.05.

**Figure 4.**
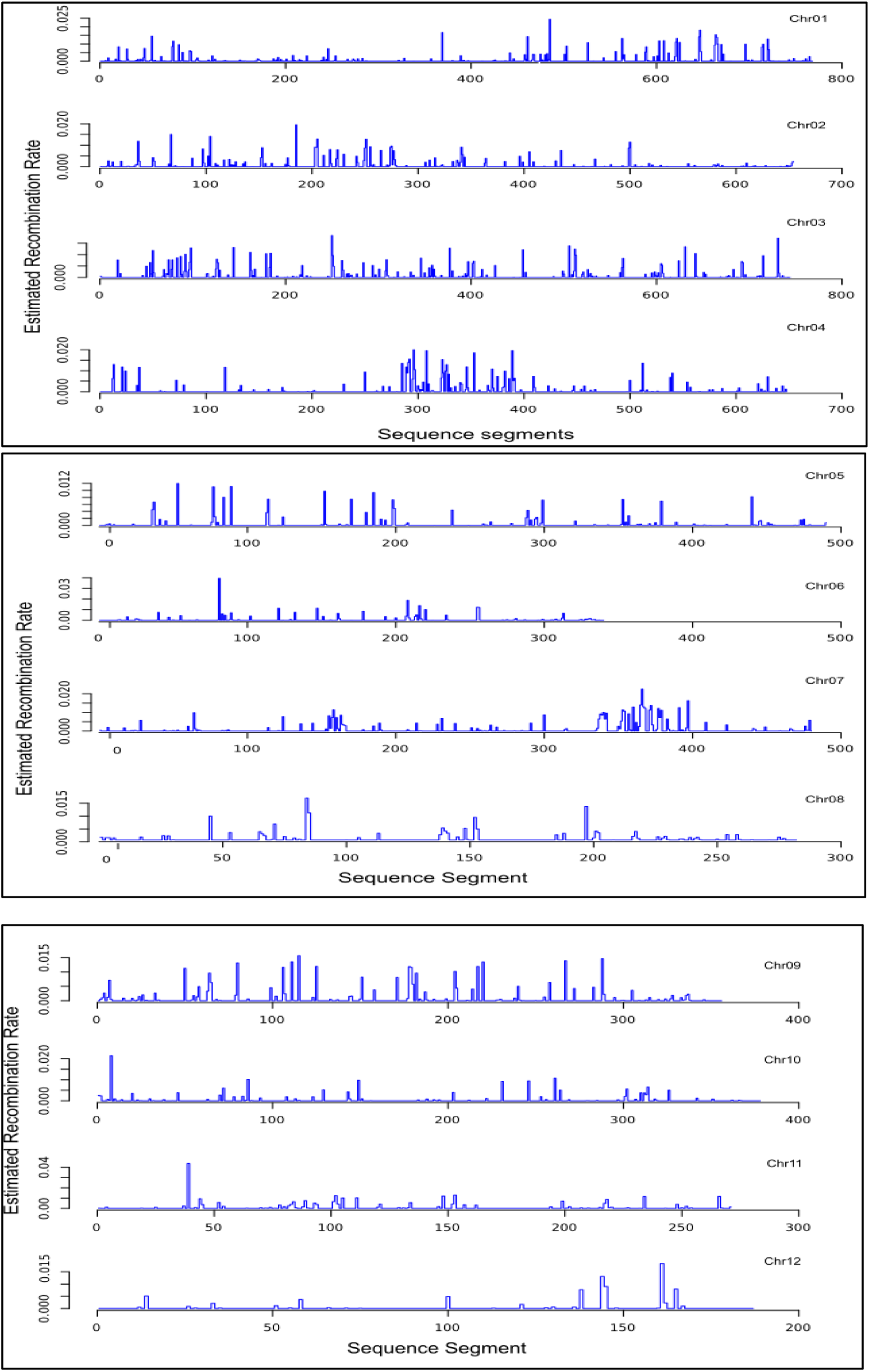
Estimated recombination map for fourteen *P. teres* isolates (10 *Ptt*, 3 *Ptm*, HR *Ptm*). Each segment has 5 kb length. The peak values indicate the higher recombination rate.

Close inspection of the aligned intergenic regions where recombination was detected revealed consistent SNP patterns in the HR *Ptm* isolate 17FRG089 (Figure 5A). In this isolate, the sequence comprised an admixed mosaic of SNPs characteristic of either *Ptt* or *Ptm* haplotypes; the observed SNP patterns spanned short genomic regions often less than 1 kb before switching between *Ptt* and *Ptm*. An example of this can be seen in the conserved intergenic region of Chr06, where *CYP51A* gene resides (Figure 5A).

**Figure 5.**
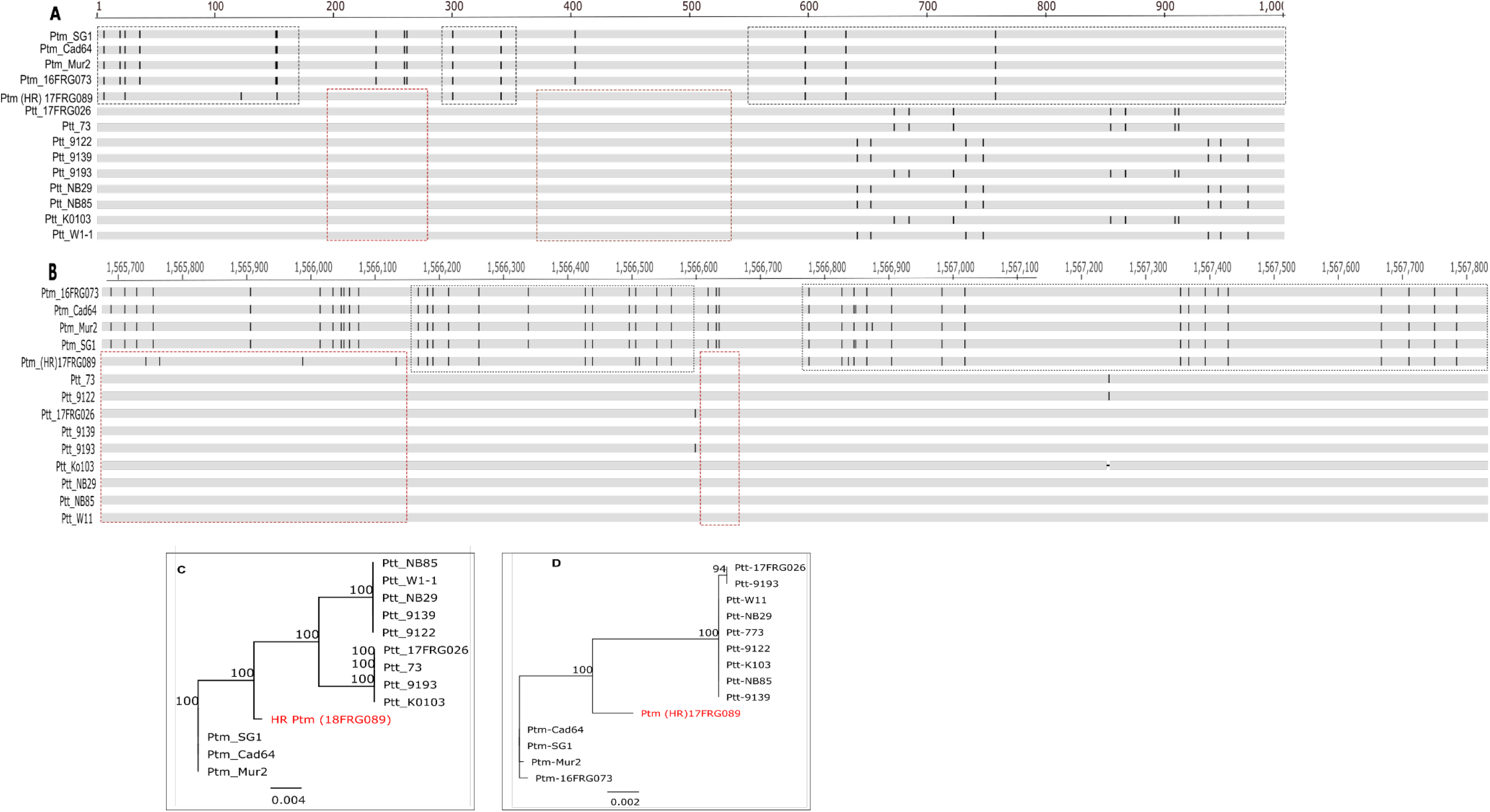
The mutation pattern observed in the HR *Ptm* isolate 17FRG089 in the intergenic region on Chr06 **(A)**, partial alignment of Chr07 **(B)** and the corresponding Neighbour-Joining trees (C, D) constructed using Juke-Cantor genetic distance model with bootstrap sampling of 1000 replicates. Vertical lines in the sequence alignment show mismatch positions. Red and black broken boxes indicate the alternating SNP pattern of HR *Ptm* isolate 17FRG089 between *Ptt* and *Ptm*, respectively.

Likewise, in a 2.1 kb region from a larger alignment of Chr07 (1,565,695 – 1,567,791), conserved HR *Ptm* SNPs were located in short segments alternating between *Ptt* and *Ptm* sequence, with a larger proportion of SNPs corresponding to *Ptm* (Figure 5B). A short 573 nucleotide segment (1,565,695 – 1,566,268) contained 20 SNPs of which the first eleven consecutive ones were mapped to *Ptm* and the last five consecutive SNPs were mapped to *Ptt*. The remaining four SNPs did not exist in either *Ptm* or *Ptt*. Five more HR *Ptm* SNPs found in an adjacent 167 nucleotide segment (positions 1,566,345, 1,566,434, 1,566,446, 1,566,504 and 1,566,512) were unambiguously mapped to *Ptt*, *Ptm*, *Ptm*, *Ptt* and *Ptm* in that order. A similar pattern could be observed in further SNP positions between (1,566,513 and 1,567,791), with three consecutive SNPs are aligned to *Ptm*, followed by another three SNPs that aligned to *Ptt* and final fifteen SNPs mapped to *Ptm*.

Neighbour-joined phylogenetic trees constructed from subsets of the intergenic region showed deviation of the HR *Ptm* isolate 17FRG089 from other *Ptm* isolates with 100% bootstrap support, but more related to *Ptm* than *Ptt* (Figure 5 C and D). The chromosome-level phylogenetic analyses is also consistent with the phylogenies constructed from the intergenic sequences. The HR *Ptm* isolate 17FRG089 is closely related to *Ptm* with the branching pattern identical on six of the chromosomes (1 – 2 and 9 – 12) and basal to the *Ptm* isolates, indicating distinct genotypic composition of the HR *Ptm* isolates (Figure 7). Interestingly, on six of the chromosomes (Chr03 – Chr08), the HR *Ptm* isolate 17FRG089 is closely related to one of the DMI sensitive *Ptm* isolates (Mur2), with a high number of SNPs shared between these two isolates indicating a common ancestral background. However, *Ptm* isolate Mur2 lacks the alternating SNP signatures observed in HR *Ptm* isolate 17FRG089.

**Figure 7.**
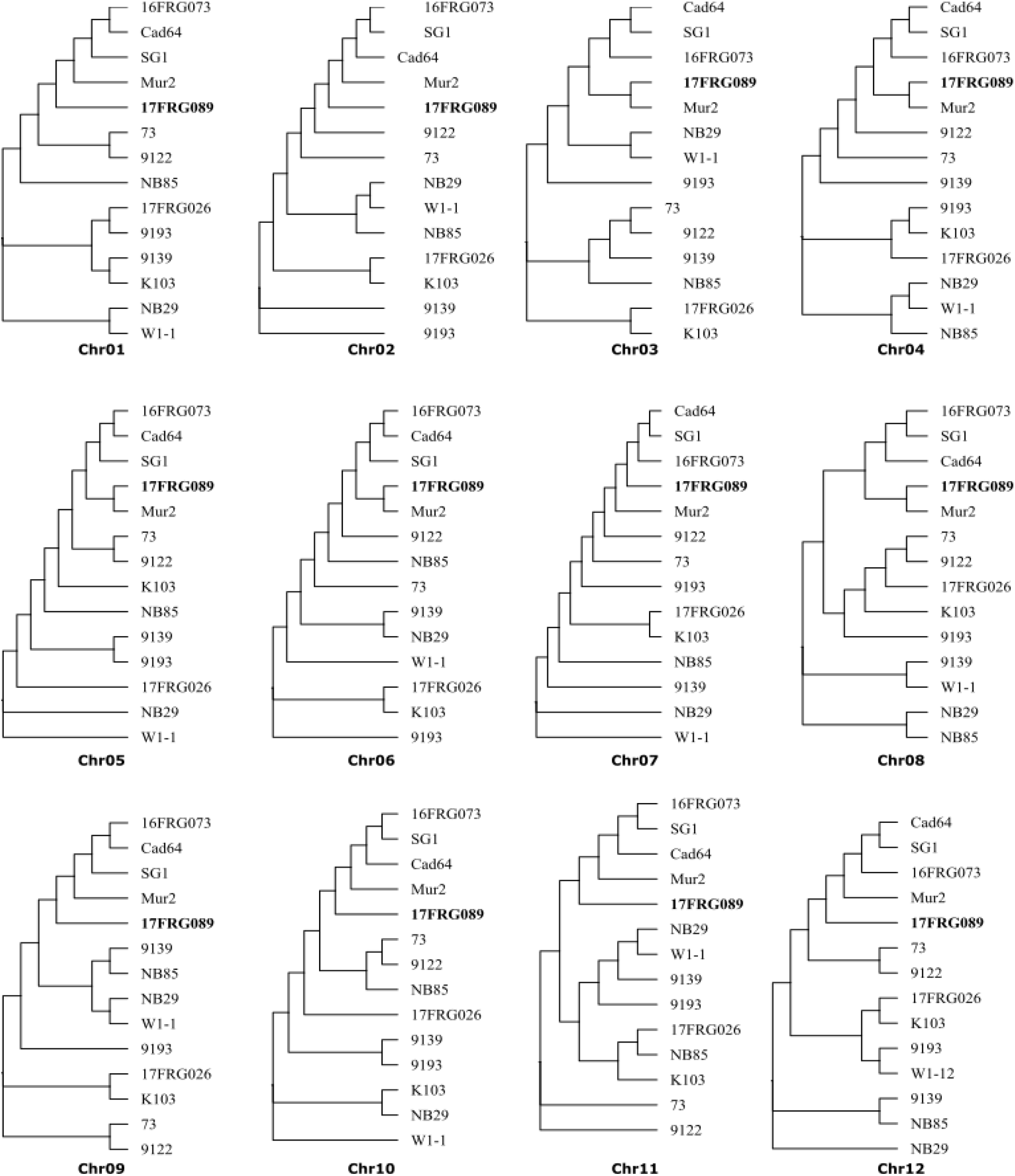
Chromosome based phylogenetic analyses of *Ptt, Ptm* and HR *Ptm* isolates. The HR *Ptm* isolate 17FRG089 is represented in bold font.

### Genetic structure analyses reveals clonal population of HR Ptm distinct from both Ptm and Ptt

The use of silicoDArT marker analyses to determine the genetic structure of the HR *Ptm* isolates, allowed to distinctly group *Ptt*, *Ptm,* barley grass *P. teres* and HR *Ptm* isolates (Figure 8). The HR *Ptm* did not cluster with *Ptt* or barley grass *P. teres*; they also did not cluster with the *Ptm* isolates but were more closely related to them. A high degree of clonality was observed among the 48 HR *Ptm*, with all of the isolates across the nine sites, separated in some cases by more than 440 km, being genetically identical to one another.

**Figure 8.**
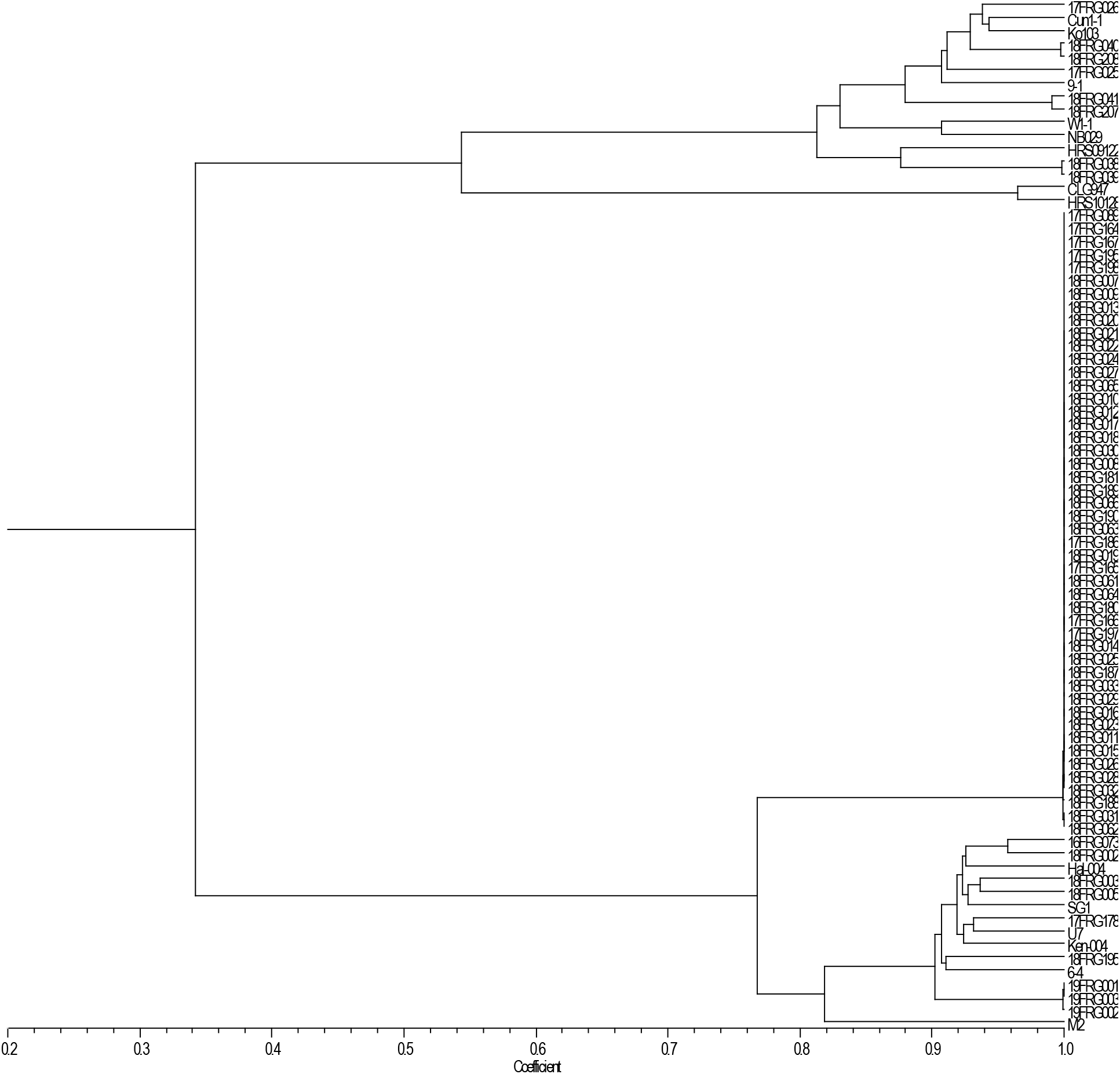
Genetic relationship among *Ptt*, *Ptm*, HR *Ptm* and barley grass *P teres*

Average Nei’s genetic distance computed among each cluster using DArT SNP showed the highest genetic distance between the HR *Ptm* and *Ptt* (0.807) isolates, while the lowest genetic distance was found between *Ptm* and the HR *Ptm* isolates (0.189). Similarly, the largest (0.981) and lowest (0.842) genetic differentiation (*F_ST_)* was observed between HR *Ptm* and barley grass *P. teres,* and between *Ptm* and HR *Ptm* (Table 3), respectively, indicating HR *Ptm* are more closely related to *Ptm*. Further molecular analyses of variance using Nei’s genetic distance also revealed significant differences in the extent of genetic differentiation among the observed clusters accounting for over 99% of the total variation (Table 4).

**Table 3.**
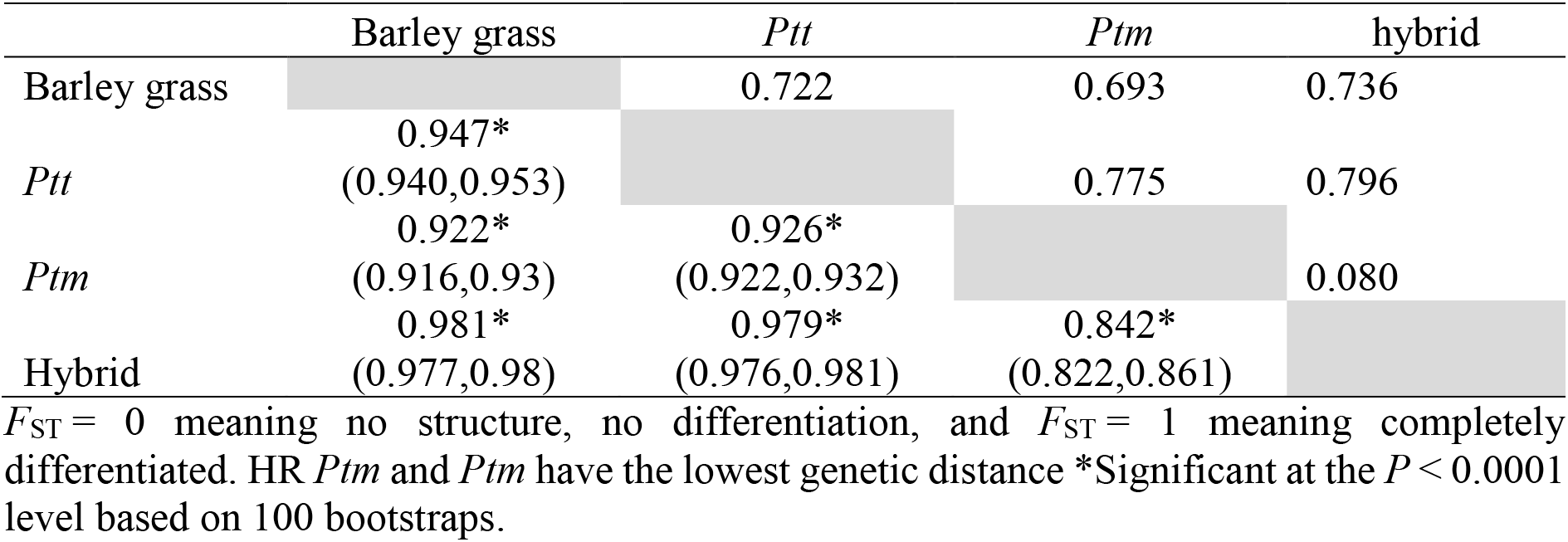
Pairwise *F_ST_* values (lower metrics) and Jaccard genetic distances (upper metrics) between four populations calculated from DArT SNP loci using 100 bootstrap. For *F_ST_*, lower and upper bound confidence intervals calculated from bootstrapping are listed within brackets.

|  | Barley grass | *Ptt* | *Ptm* | hybrid |
| --- | --- | --- | --- | --- |
| Barley grass |  | 0.722 | 0.693 | 0.736 |
| *Ptt* | 0.947*<br>(0.940,0.953) |  | 0.775 | 0.796 |
| *Ptm* | 0.922*<br>(0.916,0.93) | 0.926*<br>(0.922,0.932) |  | 0.080 |
| Hybrid | 0.981*<br>(0.977,0.98) | 0.979*<br>(0.976,0.981) | 0.842*<br>(0.822,0.861) |  |

$F_{ST} = 0$ meaning no structure, no differentiation, and $F_{ST} = 1$ meaning completely differentiated. HR *Ptm* and *Ptm* have the lowest genetic distance \*Significant at the $P < 0.0001$ level based on 100 bootstraps.

**Table 4.** Molecular analyses of variance (MANOVA) components on metrics of genetic distance between ‘Populations’

|  | df | SSD | MSD | %variance | *P*<br>value | Phi-<br>statistics |
| --- | --- | --- | --- | --- | --- | --- |
| Population | 3 | 3.024 | 1.008 | 99.985 | 0.0 | 0.99898 |
| Error | 48 | 0.007 | 0.000 | 0.015 | - | - |
| Total | 51 | 3.031 | 0.059 | 100.000 | - | - |

PCoA analyses of DArT SNP showed that 55.8%, 20.9% and 11.7% of the total variations were explained by the first, second and third PCoA observed among *Ptt*, HR *Ptm, Ptm* and barley grass *P. teres* isolates, respectively (Figure 8A). The first PCoA clearly separated HR *Ptm* from *Ptt* and barley grass *P. teres.* Similarly, the second PCoA separated barley grass *P. teres* from *Ptt*, while the third PCoA distinguished barley grass *P. teres* from *Ptm* isolates. Consistent with cluster analysis, all 48 HR *Ptm* isolates were tightly clustered on the PCoA indicating little allele frequencies overlap between HR *Ptm* isolates and other *Ptm, Ptt* or barley grass *P. teres* isolates (Figure 8A). Further analyses of SNP generated from the whole genome alignment confirmed similar result. Majority of the genetic variations were captured in the first PCoA (79%) and separated *Ptt* from the rest while the second PCoA explained only 4% of the total variation (Figure 8B). Overall, genetic variation analyses indicated that the HR *Ptm* isolates collected from WA are genetically different from other *Ptm*, and appears clonally propagated across the collection sites.

## DISCUSSION

The recent availability of genome sequences coupled with sensitive and reliable molecular detection tools have facilitated the discovery of natural interspecies hybridization between the two economically important pathogens of barley, *P. teres* f. *teres* and *P. teres* f. *maculata*. Here we employed comprehensive molecular and population genetic analyses to characterize a recently emerged population of *P. teres* isolates (HR *Ptm*) highly resistant to DMI fungicides able to induce symptoms shared by both *P. teres* f. *teres* and *P. teres* f. *maculata* characteristics.

The amplification in all the HR *Ptm* isolates of one *Ptt*-specific marker (PttQ4) provided the first indication that the HR *Ptm* isolates are hybrids. These markers were specifically developed to distinguish *Ptt* from *Ptm* and to identify hybrids between the two forms (Poudel *et al.,* 2017). Recent molecular characterization using these marker set of over 300 Australian *P. teres* isolates collected across four decades, was unable to detect any hybrids (Poudel *et al.*, 2019a). Poudel *et al.* (2019a) was also unable to detect hybrids among over 800 conidia and 200 ascospores screened from three artifical field inoculation experiments of *Ptt* and *Ptm* conducted across three seasons. A previous study by Poudel et al. (2017) using the same marker set confirmed a single field hybrid among over 200 Australian *P. teres* isolates tested. Rau *et al.* (2007) concluded that *Ptt* and *Ptm* are reproductively isolated and hybridisation between the two forms under field conditions to be either rare or absent, and furthermore that the two *formae speciales* are in an advanced stage of speciation and should be considered as distinct species. In contrast to these studies, we have found a relatively large number of hybrids isolates in the WA *P. teres* population using the form-specific markers, although samples were not collected randomly and so the true frequency of hybrids in the WA *P. teres* population cannot be estimated confidently. Given the lack of genetic diversity among the HR *Ptm* isolates, it is likely that they all derive from a single hybridisation event, in which case our findings would still accord with those from previous studies that suggest that such events are a rare occurrence in nature. A range of reproductive barriers have been hypothesized to explain the infrequent observation of *Ptt* and *Ptm* hybrids in nature (Poudel *et al.*, 2019a), including genetic incompatibilities causing unfit hybrid progeny, or reduced fitness from intermediate traits. On the other hand, Campbell and Crous (2003) found that laboratory *Ptt* and *Ptm* hybrids remained fertile, virulent and genetically stable. It is possible that the acquisition of a DMI resistance mutation in an evolutionary landscape dominated by heavy fungicide selection is compensatory to the fitness penalties that may inhere from interspecies hybridisation.

### Intergenic recombination

All the HR *Ptm* isolates carry the non-synonymous point mutation c1467a in the *Cyp51A* gene, which results in the amino acid substitution F489L in CYP51A. An identical SNP for the F489L mutation has previously been reported in *Ptt*, where it is associated with reduced sensitivity to DMI fungicides(Mair *et al.*, 2016). We evaluated whether the c1467a mutation occurred *de novo* in HR *Ptm* or was acquired from *Ptt* through genetic recombination between the two forms. Two independent genetic recombination analyses, using LDJump and DNA sequence polymorphism analyses tool (DnaSP) (Librado & Rozas, 2009) both indicated intragenic recombination within the *Cyp51A* coding sequence between *Ptt* and *Ptm*, and one of the sites involved in recombination included mutation c1467a. Previous reports have shown that intragenic recombination is an important evolutionary process in populations under fungicide selection, acting as a potential source of novel fungicide resistance alleles (Brunner *et al.*, 2008). Intragenic recombination in the *Cyp51* gene has been previously reported in *Z. tritici* (Brunner *et al.*, 2008; Estep *et al.*, 2015), and has been associated with increased resistance to DMI fungicides. The mutations in the *Cyp51* gene in European *Z. tritici* all emerged only once or twice before spreading into the broader population via gene flow, recombining to form novel combinations of resistance mutations (Brunner *et al.*, 2008). In a North American population of *Z. tritici*, the G460D mutation of *Cyp51* likely emerged *de novo* only once, before entering the broader population via at least two distinct intragenic recombination events (Estep *et al.*, 2015). Similarly, intragenic recombination has also been detected in the *Cyp51A* gene of *Rhynchosporium commune* (Brunner *et al.*, 2016). The current study is, to our knowledge, the first report of interspecific recombination involving a gene associated with fungicide resistance in *P. teres* species.

The intergenic and whole chromosome recombination analyses using *de novo* assembled genome sequences of each *P. teres* isolates provided strong evidence of genetic exchange between the two forms, consistent with analyses using form-specific markers and recombination analyses of the *Cyp51A* gene. Previous studies have indicated that *de novo* assembly-based analyses enables identification of highly variable genomic regions involved in hybridization which are missed through raw read mapping approach (Feurtey *et al.*, 2019). Studies of *Zymoseptoria ardabiliae* and *Z. tritici* indicated high recombination rates in intergenic regions (Stukenbrock & Dutheil, 2018). Similar evidence of recent hybridization has been reported within several species of the *Zymoseptoria genus* (Feurtey *et al.*, 2019). Recombination has also been detected in intergenic regions between members of the *Histoplasma* genus (Maxwell *et al.*, 2018).

### Population structure

The neighbour joining phylogenetic analyses using subset of intergenic sequences showed that the HR *Ptm* isolates (represented by 17FRG089) lie between *Ptt* and *Ptm* with 100% bootstrap support. Further chromosome-level phylogenetic tree analyses were also concordant to phylogenetic trees generated from the intergenic sequences, indicating that the HR *Ptm* isolates are genetically distinguishable from both *Ptt* and *Ptm* isolates. The whole genome-based PCoA also separated HR *Ptm* (17FRG089) as distinct from *Ptt* and *Ptm*, suggesting unique genetic constitution of the HR *Ptm* isolate in the current study. Since the HR *Ptm* was represented with single genome sequence, we utilised DArT SNP analyses with a larger number of HR *Ptm* isolates to capture the genetic variation that existed among *Ptt, Ptm*, HR *Ptm* and *P. teres* isolates from barley grasses. The smallest computed genetic distance (0.080) was between HR *Ptm* and other *Ptm* isolates, suggesting that the genetic makeup of the HR *Ptm* is mainly *Ptm,* likely resultant from multiple backcrossing of the progenies following the hybridisation event. The clustering of the HR *Ptm* into a single group on dendrogram and PCoA (Figure 8 and 9) showed an absence of genetic diversity among the HR *Ptm* isolates. The HR *Ptm* were sampled from sites separated by up to 440 km, from Frankland River in the Great Southern region to Gibson in the Esperance region in WA, suggesting a clonal expansion of the HR *Ptm* isolates as a potential explanation for their spread across a wide geographical area in a very short period of time. These results are congruent with asexual selection of progeny harbouring DMI-resistance alleles under the selective pressure of DMI use, following a single hybridisation event. DMI-resistant *A. fumigatus* isolates from India were highly related despite their collection points were more than 1000 km apart (Abdolrasouli *et al.*, 2015). The authors concluded that the DMI-resistance mutations were therefore likely associated with an *A. fumigatus* highly fit genotype that had undergone a recent, rapid selective sweep. A similar dynamic may account for the lack of observed variation among isolates of the HR *Ptm* genotype, with DMI fungicide use or possibly varietal choice acting as the selective sweep within the WA *Ptm* population.

**Figure 9.**
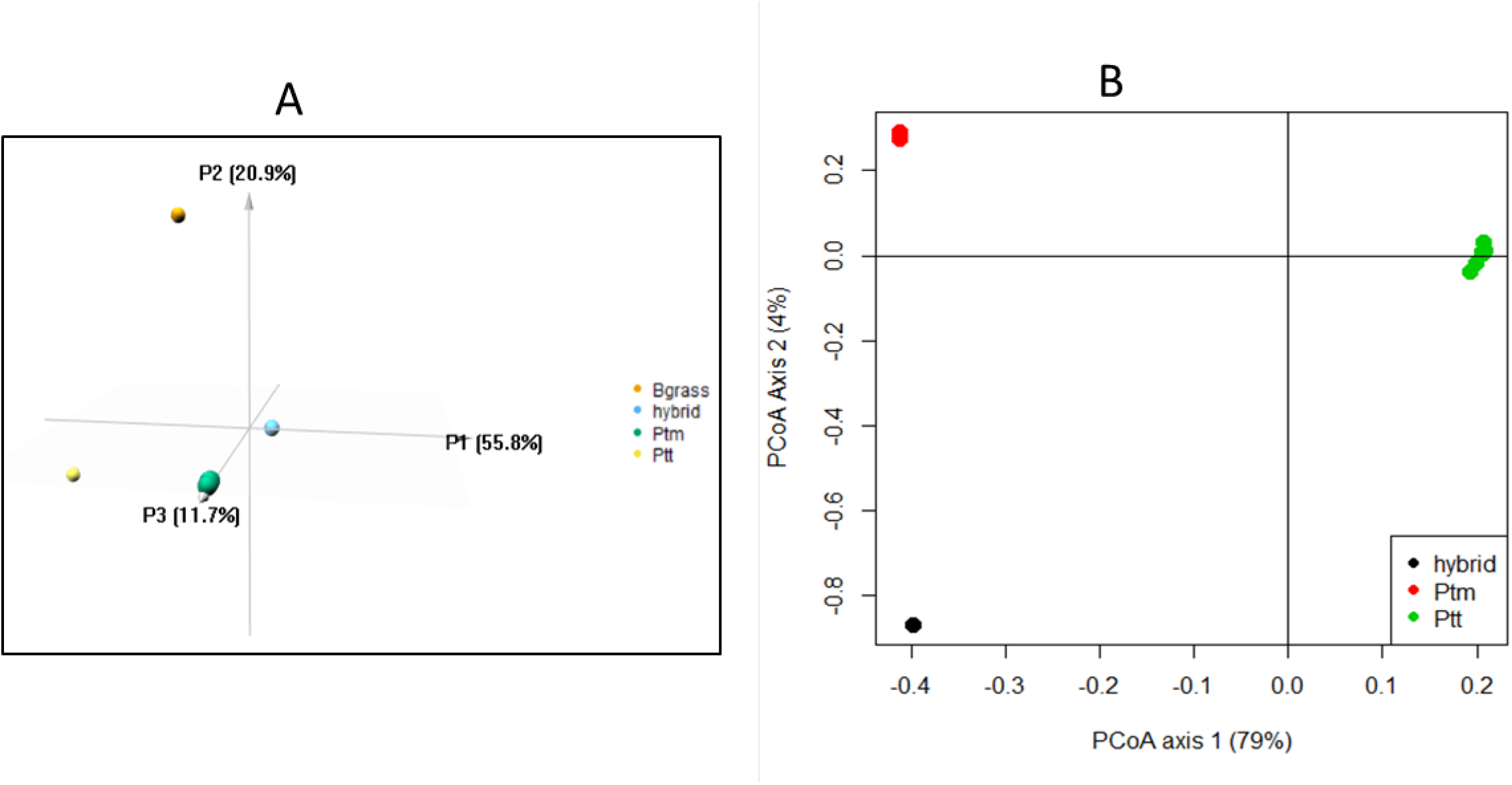
Distribution of *Ptt, Ptm*, HR *Ptm* and barley grass *P. teres* isolates in different quadrants based on principal coordinate analyses using SNP DArT markers **(A)** and whole genome SNP data **(B)**. **(A)** In SNP DArT marker analysis, PCoA 1, PCoA 2 and PCoA 3 explain 55.8% and 20.9% 11.7% of the total variations, respectively**. (B)** In Large scale PCoA analyses using SNP generated from genome, the first PCoA alone explained 79% of the total variation.

### Implications

Campbell and Crous (2003) previously raised the epidemiological implications of natural hybridisation between *Ptt* and *Ptm*, with the possibility of new genotypes thereby being introduced into populations. Interspecies recombination is increasingly acknowledged for its significant contribution to pathogen diversification (Feurtey & Stukenbrock, 2018). Genetic hybridization studies between the grass symbionts *Epichloe festucae* and *Epichloe gansuensis* indicated that genetic introgression of fungi living on the same hosts significantly contribute to their adaptive evolution (Zhang *et al.*, 2018). In some cases hybridisation can even lead to the formation of novel species, as in the wheat pathogen *Zymoseptoria pseudotritici* (Stukenbrock *et al.*, 2012), or novel *formae speciales*, as in the triticale pathogen *Blumeria graminis* f. sp. *triticale* (Menardo *et al.*, 2016). Although true population frequencies cannot be estimated confidently, it is apparent that among the DMI-resistant WA *Ptm* isolates found so far, the clonal HR *Ptm* genotype was clearly predominant (Mair *et al.*, 2020), suggesting the possibility of lineage replacement.

Historically, *Ptm* was reported as much less prevalent than *Ptt* in WA; a 1995-96 survey was the first to observe the spot form net blotch disease outside of the northern barley-growing region of WA, while it reached epidemic levels in the southern agricultural regions of WA in 1997-1998 (Gupta & Loughman, 2001). In subsequent years, *Ptm* became much more prevalent across WA southern barley-growing regions, aided by the wide spread adoption of susceptible varieties such as Gairdner, Hamelin, Vlamingh and Baudin (Gupta *et al.*, 2012). Campbell *et al.* (2002) suggested that growing barley cultivars susceptible to both forms in very close proximity potentially favour sexual reproduction between the two forms. The lack of resistant cultivars together with the introduction of stubble retention practices and high inclusion of continuous barley, has led to an increase in disease incidence and severity in recent years (Gupta *et al.*, 2012)(McLean *et al.*, 2009). These conditions have likely increased the opportunities for hybridisation between the two pathogens as they co-exist infecting the same host. The HR *Ptm* isolates were mainly associated with barley cv. Oxford, a variety that has become increasingly susceptible to *Ptt* over recent years with the spread of the new, aggressive ‘Oxford virulent’ pathotype, particularly around the Esperance and Great Southern growing regions (Shackley, 2019). The other varieties with which the HR *Ptm* genotype was associated (albeit at lower frequencies), Planet and La Trobe, are also closely related to Oxford.

The use of DMI fungicides in barley in Australia begun in 1995 and has increased substantially in subsequent decades (Tucker *et al.*, 2015). It appears likely that the *Ptt-Ptm* hybridization event took place in the last decade, because the point mutation c1467a in *Cyp51A* gene is absent in *Ptm* collections prior to 2017 (Mair *et al.*, 2020), and the same *Cyp51A* point mutation in *Ptt* was found only from 2013 onwards (Mair *et al.*, 2016). Analyses of samples from older collections may allow us to more accurately estimate the exact time of hybridisation using a population genomics approach.

*Ptm* conidia are generally understood to be dispersed primarily by wind and rain splash over relatively short distances (Poudel, 2018), so other factors, such as dispersal on hay or possibly on seed, must account for the observed wide geographical distribution of the HR *Ptm* genotype in WA in a relatively short period of time. Although only *Ptt* has been reported to be seed transmitted (McLean *et al.*, 2009), it is not clear if this would also be true for any putative *Ptt-Ptm* hybrids, and it is possible that genes involved in *Ptt* seed transmission have introgressed during hybridisation. Additional genes affecting virulence on different host varieties may also have been acquired by the hybrid, which may help account for the observed preponderance of HR *Ptm* isolates being derived from cv. Oxford and a small number of closely-related cultivars. In light of these findings, there is an urgent need for the deployment of integrated disease and resistance management strategies that take into account not just the effect of fungicide use and varietal choice in both *Ptt* and *Ptm*, but also the possible exchange of fungicide resistance and virulence genes between the two forms.

## ACKNOWLEDGEMENT

The authors would like to thank the sample contributions made by growers and agronomists. This study was conducted by the Centre for Crop and Disease Management, a joint initiative of Curtin University and the Grains Research and Development Corporation (research grant CUR00023). We also acknowledge the assistance of resources from the National Computational Infrastructure (NCI Australia), an NCRIS enabled capability supported by the Australian Government.

